# Machine-learning model led design to experimentally test species thermal limits: the case of kissing bugs (Triatominae)

**DOI:** 10.1101/2020.10.05.326017

**Authors:** Jorge E. Rabinovich, Agustín Alvarez Costa, Ignacio Muñoz, Pablo E. Schilman, Nicholas Fountain-Jones

## Abstract

Species Distribution Modelling (SDM) determines habitat suitability of a species across geographic areas using macro-climatic variables; however, micro-habitats can buffer or exacerbate the influence of macro-climatic variables, requiring links between physiology and species persistence. Experimental approaches linking species physiology to micro-climate are complex, time consuming and expensive. E.g., what combination of exposure time and temperature is important for a species thermal tolerance is difficult to judge *a priori*. We tackled this problem using an active learning approach that utilized machine learning methods to guide thermal tolerance experimental design for three kissing-bug species (Hemiptera: Reduviidae: Triatominae), vectors of the parasite causing Chagas disease. As with other pathogen vectors, triatomines are well known to utilize micro-habitats and the associated shift in microclimate to enhance survival. Using a limited literature-collected dataset, our approach showed that temperature followed by exposure time were the strongest predictors of mortality; species played a minor role, and life stage was the least important. Further, we identified complex but biologically plausible nonlinear interactions between temperature and exposure time in shaping mortality, together setting the potential thermal limits of triatomines. The results from this data led to the design of new experiments with laboratory results that produced novel insights of the effects of temperature and exposure for the triatomines. These results, in turn, can be used to better model micro-climatic envelope for the species. Here we demonstrate the power of an active learning approach to explore experimental space to design laboratory studies testing species thermal limits. Our analytical pipeline can be easily adapted to other systems and we provide code to allow practitioners to perform similar analyses. Not only does our approach have the potential to save time and money: it can also increase our understanding of the links between species physiology and climate, a topic of increasing ecological importance.

**Author summary:** Species Distribution Modelling determines habitat suitability of a species across geographic areas using macro-climatic variables; however, micro-habitats can buffer or exacerbate the influence of macro-climatic variables, requiring links between physiology and species persistence. We tackled the problem of the combination of exposure time and temperature (a combination difficult to judge *a priori*) in determining species thermal tolerance, using an active learning approach that utilized machine learning methods to guide thermal tolerance experimental design for three kissing-bug species, vectors of the parasite causing Chagas disease. These bugs are found in micro-habitats with associated shifts in microclimate to enhance survival. Using a limited literature-collected dataset, we showed that temperature followed by exposure time were the strongest predictors of mortality, that species played a minor role, that life stage was the least important, and a complex nonlinear interaction between temperature and exposure time in shaping mortality of kissing bugs. These results led to the design of new laboratory experiments to assess the effects of temperature and exposure for the triatomines. These results can be used to better model micro-climatic envelope for species. Our active learning approach to explore experimental space to design laboratory studies can also be applied to other environmental conditions or species.

## Introduction

The main environmental requirements for any organism to be in thermodynamic equilibrium over a reasonable length of time in order to survive are well known for more than a half century [1]. Of these, radiation absorbed, wind speed, and air temperature are physiological requirements for a certain body temperature or temperature range, and are referred to as the ‘climate space’, and constitute the conditions which animals must fulfil in order to survive [2]. Much effort has gone into applying Species Distribution Modelling (SDM) to model climate space across different geographic areas to determine habitat suitability, including species that are disease vectors [3–9]. In general, the SDM methodology utilizes macro-climatic variables to predict the distribution of a species; however, for many disease vector species natural and human-made micro-habitats can buffer or exacerbate the influence of the macro-climatic variables [10,11]. While these models increasingly harness high resolution climatic data down to ~1km^2^ (e.g., WorldClim 2 [12]), the links between physiology, disease transmission and vector survival to micro-climate are opaque. Consequently, the use of these climatic variables leads to some caveats on the reliability of the suitability estimates produced by the SDM. However, accurately quantifying micro-climatic relationships and tolerances pose significant problems for most disease vector species.

Mathematical models have helped predict operative temperatures but also show how evaporative cooling and metabolic heating might cause the body temperature of an organism to deviate from the operative temperature [13]. However, despite mathematical models have provided important insights into what factors influence operative temperatures, they are impractical for mapping thermal environments at a sufficient resolution to understand selective pressures on behavior and physiology [14,15]. This is because to compute operative temperatures from a mathematical model many variables must be known (solar radiation, ground reflectivity, air temperature, ground temperature, and wind speed), which represents an overwhelming task for a large number of locations as needed for fine-scale mapping [15]. Operative temperatures have been computed for a limited number of microclimates, such as full sun and full shade, and for a few animal species [16–18], but a general methodology is still needed.

Because of their use of a variety of micro-habitats, the trouble of using macroclimatic variables to predict vector species distributions is particularly serious [19]. This is true for the kissing bugs (Hemiptera: Reduviidae: Triatominae), a group of species vectors of *Trypanosoma cruzi*, the parasite causing Chagas disease in Latin America. This disease is endemic in 21 countries, and it is estimated that affects between 6 and 8 million individuals (with 25 million people at risk of infection), resulting in approximately 12,000 deaths per year [20,21]. The triatomines are a subfamily that comprise around 150 species grouped in 18 genera and six tribes [22]. A commonly observed behavior of most of these species is to enter domestic and peri-domestic structures (“intrusion”), with some of them trying to colonize the human habitat (“domiciliation”), making the sylvatic species a possible source of infection by *T. cruzi* [23]. Due to their generally nocturnal habits and hidden refuges, they are not only inconspicuous and hard to collect in the field, but also endure extreme values of macro-climatic variables that would not allow them to survive without the use of micro-habitats [24].

For triatomines, as with other disease vector species, there is very limited information available to adequately link insect physiology to micro-climatic variables. Much more experimental work is necessary, but those experiments are laborious, costly, and demanding a large number of insects. Additionally, the experimental design is a complex one for, in general, the micro-climatic variables (*e.g*., temperature) need to be included with at least two factors: the value of the micro-climatic variable itself, and the exposure time (or duration) to each value of the micro-climatic variable. It is difficult to guess beforehand the limits and number of those two factors, or the impact of each of them (and their combination) on the demographic parameters, in order to design a laboratory experiment.

To facilitate the design of such kind of experiments (which, in theory, could involve hundreds or thousands of combinations) we propose a methodology based upon a machine learning pipeline approach [25] in order to predict the survival of triatomines by different combinations of micro-climatic temperature values and exposure times. This approach leverages recent advance in machine learning to construct powerful but interpretable predictive models that can guide what combinations of variables could be important in shaping a species thermal limit and thus can configure feasible experimental designs. Importantly these models can quantify complex non-linear interactions between variables that can be difficult to include *a priori* in a model.

Finding computational solutions to guide experimental design or ‘active learning’ is not a new idea and has been used to guide experiments to better understand complex gene regulatory networks [26]. Active learning is an iterative process in that a model is formed from preliminary data that guides new experiments that in turn generates novel data that is used to update the original model (Fig 1). For example, machine learning approaches have been successfully used to guide gene knock-out experiments where the number of experiments is quadratic to the number of genes (see [26] for a review on the topic). However, active learning approaches are rarely applied to explore species thermal limits. We show the utility of the approach to better understand the thermal ecology of an important group of disease vector species that could easily be adapted to guide ecological experiments more broadly.

**Fig 1.**
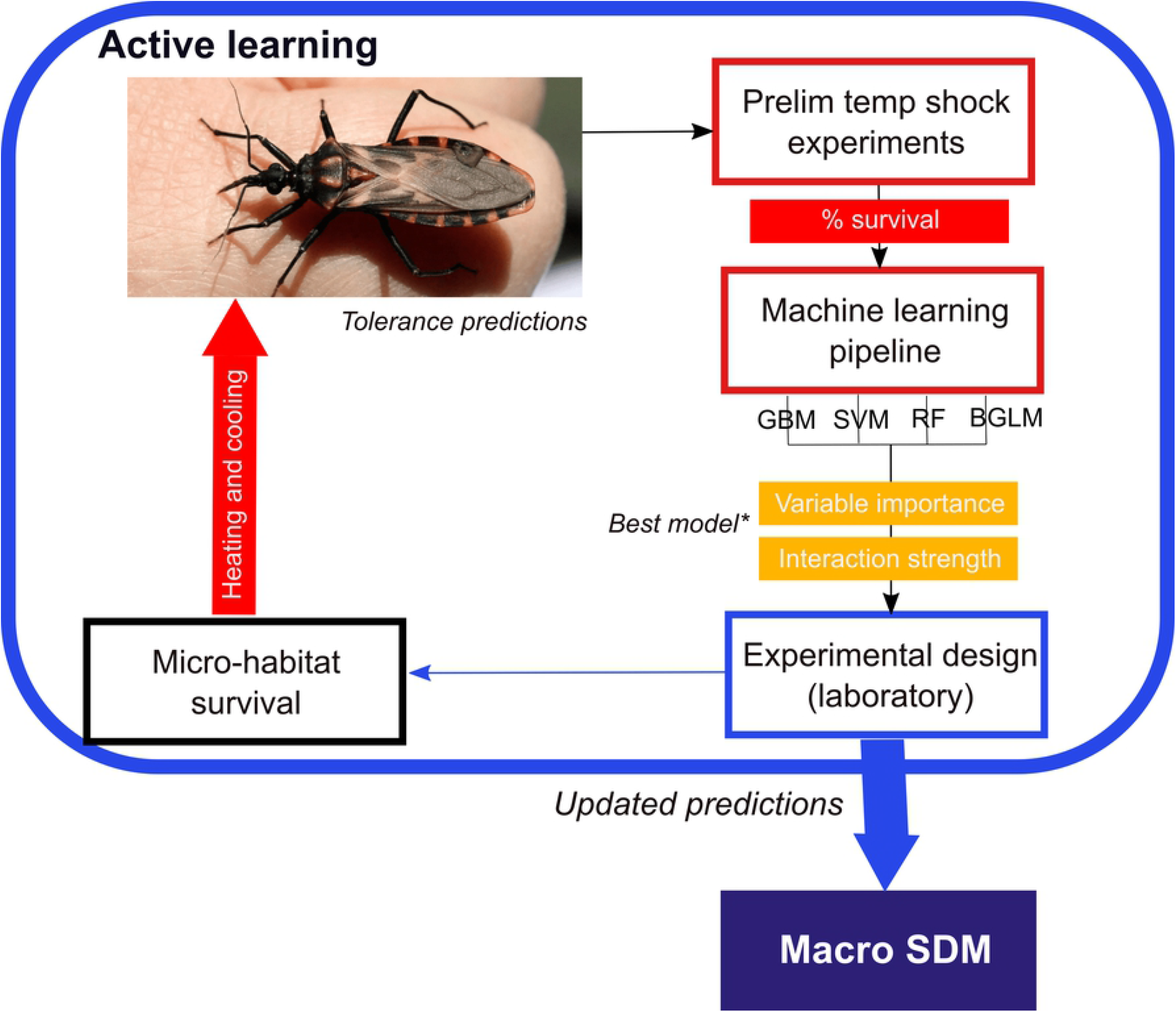
Schematic description of the active learning approach used in this study. We have highlighted the context and purpose of the work, the approach and methods, and an outline of the application of the main results. GBM: Gradient boost model, SVM: Support vector machine, RF: Random forests, BGLM: Bayesian general linear model. *Best model: model with the highest root mean square error (RMSE).

## Materials and methods

### Available laboratory information for triatomines

#### Data source

The only previous data available were the results from thermal shock experiments (usually at 40 °C) applied on different stages and adults of several triatomine species, exposed for 1 and 12 hours. In total we were able to find seven bibliographic sources with adequate data for three species of triatomines (*Triatoma infestans, Rhodnius prolixus*, and *Panstrongyus megistus*). See Data Cases below.

#### Data preparation

As in most of the cases results were presented as survival values (*l_x_*, with values from 1 to ≥0, which express the proportion of the initial number of individuals alive at day *x*, and where 1 ≤ *x* ≤ 30, 30 being the number of days of the control period of all experimental insects), we converted these survival values to mortality (*d_x_*, or number of individuals dying in day *x*, where again x is from day 1 to day 30 of the control period; we expressed the result of this conversion as *d_x_*= (*l_x_* - *l_x-1_*). The mortality values were inserted into a file with a database structure, and were considered as the end points (as percentages), for each combination of temperature, exposure time, and stage (and sex in the case of adults) predictor variables (hereafter ‘features’ in line with computer science terminology), making a total of 228 combinations (Supplementary Material 2). Table 1 represents a summary of the different cases analyzed.

**Table 1.** Source and characteristics of the original information.

| Species | Temperature | Exposure | Stage | N | Period | Source |
| --- | --- | --- | --- | --- | --- | --- |
| *Panstrongylus megistus* (D) | 28, 40 | 1, 12 | 3,4,5,Ad | 100 | 30 | [27] |
| *Panstrongylus megistus* (D) | 5, 28 | 1, 12 | 5 | 53, 100 | 30 | [28] |
| *Panstrongylus megistus* (D) | 35, 40 | 1, 12 | 5 | 53, 100 | 30 | [28] |
| *Panstrongylus megistus* (S) | 28, 40 | 1, 12 | 3,4,5,Ad | 100 | 30 | [27] |
| *Rhodnius prolixus* | 35-44 | 1, 24 | 1 | 10 | 1 | [29] |
| *Triatoma infestans* | 30, 40 | 1, 12 | 3,4,5,Ad | 50 | 30 | [30] |
| *Triatoma infestans* | 35, 45, 50, 55 | 0.17 | E,1,2,3,4,5,Ad | 30 | 45 | [31] |
| *Triatoma infestans* | 30 | 1, 12 | 5 | NA | 30 | [32] |

Summary information of the cases with data available on the effect of various temperatures on the survival of triatomines obtained from the bibliography, and fed to the machine learning algorithm. For the species *Panstrongylus megistus* D and S represent domiciliary and sylvatic origin, respectively; Temperature is in °C; Exposure is measured in hours; Stage represents the triatomine developmental stage used in the experiments (E: eggs; 1-5: first to fifth nymphs; Ad: adults); N is the number of individual replicates per temperature and exposure combination; Period: period of observation (days) after experimental exposure to determine survival. The complete dataset can be found in the Supplementary Material 2.

### The machine learning pipeline approach

We adapted the framework from [25] using our collated feature set to predict percentage mortality as a regression problem. Initially we checked for correlations between features prior to running our machine learning pipeline. We compared support vector machine, gradient boosting model (GBM), Bayesian generalized linear models as well as linear models fitted with ordinary least squares. In each case we used 10-fold cross validation to compute model performance (root mean square error, RMSE) and to prevent overfitting the model. The model with the lowest RMSE and thus highest predictive performance was further interrogated. We also compared predictive performance by calculating *R^2^*, in this case the correlation between the observed and predicted values [33]. To interpret how each feature was affecting model performance we computed ranked feature importance using model class reliance approach [34] and plotted individual responses using individual conditional expectation curves [35]. Interactions in the model were quantified by calculating Friedman’s *H* statistic [36] and visualized plotting multidimensional partial dependency plots. See [25] for more details and https://github.com/nfj1380/ThermalLimits_ActiveLearning for the code used.

## Results

### The machine learning pipeline results

The GBM model had the highest predictive power compared to the other algorithms we tested, with an *R^2^* of 0.8 and RMSE of 18.2. In this model, temperature followed by the number of hours exposed were the strongest predictors of mortality based on the preliminary experimental results (Fig 2a). Species played a more minor role in our predictive models with life stage the least important of the variables included in the model (Fig 2a). When we further interrogated our model, we not only found some strong non-linear effects of each variable on mortality (Fig 2b-e) but some striking non-linear interactions between variables (Fig 3). For example, triatomine mortality was generally stable up until the 39°C mark when it rapidly increased before plateauing between 40-44°C, and this response was an essential component of the experimental design for the combination of temperature and exposure time on insect’s survival (see below). However, mortality predictions for some individuals in the dataset were much lower than others which is evidence that interactions are important in this predictive model (indicated by the spread of the lines in Fig 1b, see [37]). The relationship between exposure and mortality was even more variable across individuals showing no clear trend (Fig 2c). Exposure time, however, had the strongest interactions with the other variables (Fig 3a), with the relationship between exposure time and temperature being of the strongest predictive importance. Low mortality was predicted with short exposure times (<7 hours to temperatures between 0-39°C), yet any exposure to temperatures above 42-44 °C was associated with high mortality, although an exact threshold was not evident. Our model predicted that temperatures between 10-39°C to be optimal for triatomine survival with no effect of exposure time on mortality. Our analysis also revealed that *T. infestans* experienced lower mortality overall (Fig 2c), and this may be linked to temperature as we found lower mortality of this species compared to the others at temperatures >39°C.

**Fig. 2.**
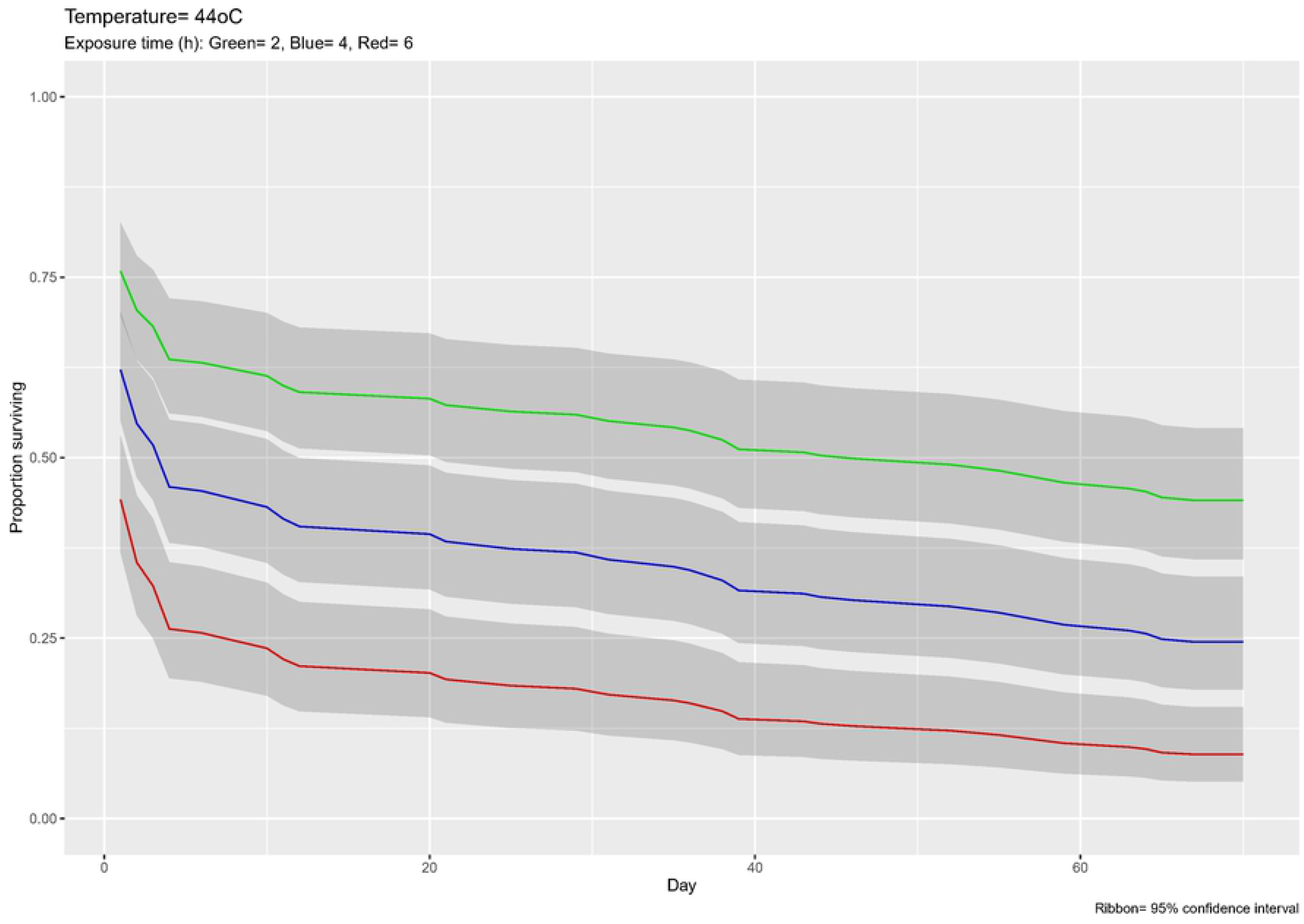
Results from the best performing machine learning model. Variable importance plot (a) and centered individual expectation plot (cICE) plots (b-e) from the best performing machine learning model (GBM). Y axes represent the marginalized effect of each variable on vector estimated mortality (%) whilst controlling for the effect. The red line represents the average effect of the variable of triatomine survival.

**Fig. 3.**
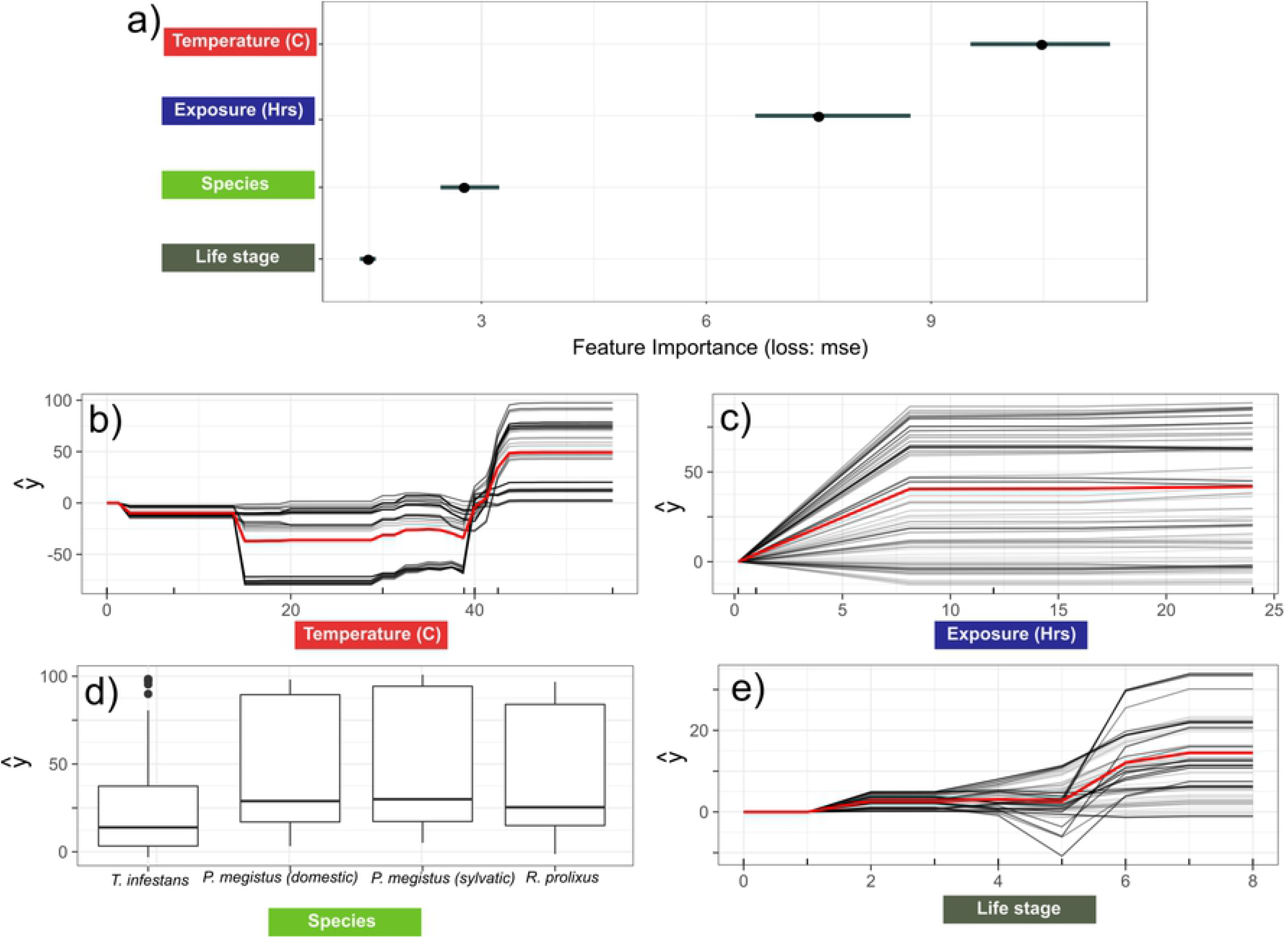
Interaction between variables as detected from the machine learning model. Plots showing the overall interactive strength of each variable (a), the top three interactions between variables based on Friedman’s *H* (b), followed by 2D partial dependencies of the most important interactions in our GBM model using Friedman’s *H* Index (c-d), where ŷ is the estimated mortality (%). Darker red in the heatmap (c) reflect higher predicted mortality. Fig 1 of the Supplementary Material 1 shows the interaction between exposure time and species.

### Machine learning led experimental design

The results of our machine learning model revealed that any exposure to temperatures above 42-44 °C was associated with high mortality. Subsequently, we exposed insects to one of three temperatures (40, 42 and 44 °C) for different exposure times (1, 2, 4, 8 and 12 hours). We used fifth-instar nymphs of *T. infestans* provided by the National Chagas Control Service (Córdoba, Argentina). Insects were fed on live chicken two weeks before experiments, and they were acclimated in the laboratory at 25 ± 0.5°C, and 12:12 light/dark photoperiod (light on 08:00 am) for one week, because thermotolerance showed a plastic response in this species [38].

The insects were placed in a chamber (PTC-1 Peltier Effect Cabinet; Sable Systems International (SSI), Las Vegas, NV, USA), connected to a temperature controller (Pelt-5 (SSI)) set at 40, 42 or 44°C for each of the exposure times. A group of insects handled in the same way as the experimental groups, but without being exposed to high temperature, and kept at 25°C, were used as the control group (C). After their exposure to the treatment the insects were maintained at the basic laboratory conditions described above, for 70 days. Ten insects were used for each replicate; the 40°C treatment was replicated twice, while the 42 and 44°C treatments were replicated three times. A total of 480 insects were used. The number of dead and molted insects were recorded each day after the heat treatment. Molted insects were eliminated from the survival analysis. A survival percentage was calculated as the ratio of number of live insects over the total number of treated insects (*i.e.*, the “initial” number) for each replicate and treatment. To determine the effects of heat for different temperatures and exposure periods on survival, a Cox analysis (using the package *survival* of the R language [39]) was performed for each experimental temperature, *i.e*., 40, 42 and 44 °C. Also, a GLM (Generalized Linear Model) analysis was carried out to determine the importance of the covariates (the treatment temperature and exposure times of the laboratory experiments), as well as their interactions, on mortality; the GLM enables the use of linear models in cases where the response variable has an error distribution that is non-normal. The GLM fit to the data was then used to predict the insects’ mortality for various combinations of temperatures and exposure times; those mortality predictions were converted into age specific survival values (*l_x_*) and then used to estimate the life expectancy (days) after the heat treatment (*e_0_*). All these calculations included the 95% confidence intervals and were carried out in R [40](R Core Team, 2020); see https://github.com/nfj1380/ThermalLimits_ActiveLearning for the code used.

### The laboratory experiment results

At 40 °C survival did not differ significantly across exposure periods (Log-rank test, χ2 = 5.33, df = 5, *p*= 0.377) (Table 2). However, survival differed significantly across exposure periods at the other two temperatures, *i.e*., 42 °C (Log-rank test, χ2 = 66.92, df = 5, *p*< 0.001), and 44 °C (Log-rank test, χ2 = 138.25, df = 5, *p*< 0.001). At 42 °C, the exposure periods of 8 and 12 h showed a lower mean survival time than the other exposure times (Table 2). At 44 °C, the mean survival time of 2 and 4 h treatments were lower than the control and 1 h treatment, and 8 and 12 h showed the lowest mean survival time from all treatments (Table 2).

**Table 2.** Laboratory experiment results of the effect of temperature on survival of 5th instar T. infestans.

| Temp. (°C) | Control | Exposure periods (h) |  |  |  |  |
| --- | --- | --- | --- | --- | --- | --- |
|  |  | 1 | 2 | 4 | 8 | 12 |
| 40 | 64.1 ± 4.3a | 67.0 ± nda | 65.6 ± 3.6a | 69.5 ± nda | >70 a | 64.7 ± 5.1a |
| 42 | 65.9 ± 2.2a | 64.6 ± 3.1a | 54.6 ± 5.4a | 63.8 ± 4.3a | 37.5 ± 5.8b | 20.83 ± 5.5b |
| 44 | 68.8 ± 1.2a | 22.2 ± 5.3a | 21.97 ± 5.9b | 13.9 ± 4.4b | 1 ± 0c | 1 ± 0c |

Survival time (mean ± SE) in days of fifth nymphs of *T. infestans* under different exposure periods (h) and temperatures (Temp, °C). Different letters indicate significant differences between the exposure times of each temperature. nd: the parameter could not be estimated.

The fit of the mortality data using the survival function (with interaction) of the Cox analysis showed that both covariates (temperature and exposure times) were highly significant, as well as their interaction, in determining the mortality level (Concordance= 0.872 (se = 0.015), Likelihood ratio test= 295.9 on 3 df, *p*≤ 2e-16, Wald test = 276 on 3 df, *p*≤ 2e-16, and Score (logrank) test = 447.6 on 3 df, *p*≤ 2e-16). Fig 4 shows the result of the survival curve estimation for only one temperature (44 °C) and three exposure times (2, 4, and 6 hours). The survival curves for all combinations of temperature and exposure times are given in Section 2 of the Electronic Supplementary Material 1.

**Fig 4.**
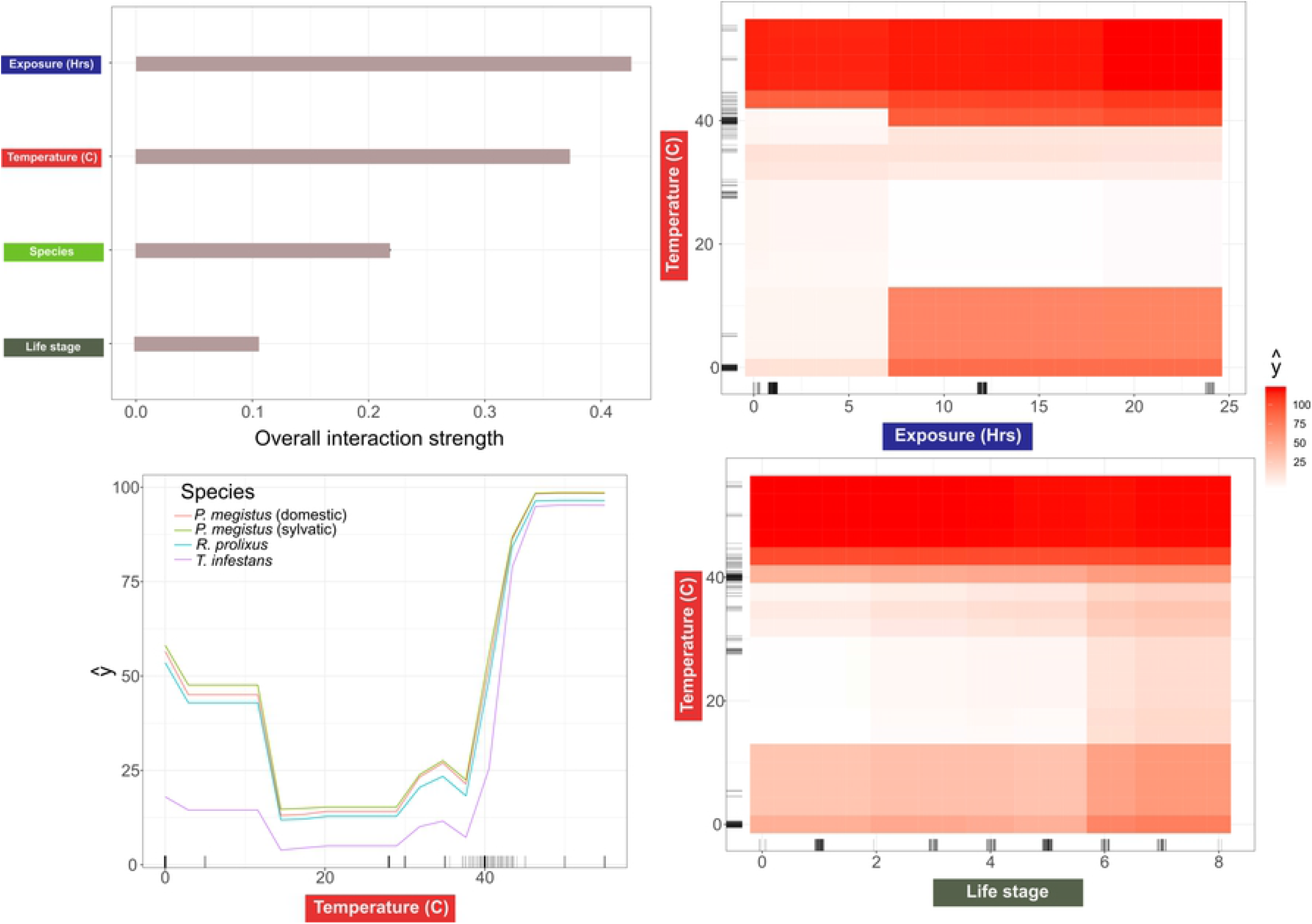
Results of Cox analysis of laboratory experiments. The survival curves (*l_x_*) predicted by the Cox analysis for 44 °C temperature and three exposure times (2, 4, and 6 hours).

The GLM analysis confirmed this kind of results, except that it showed a much weaker effect of temperature (NS at the 5% level: *p*= 0.0552) and a highly significant effect of exposure time and the interaction of temperature with exposure time (*p*= 0.00408, and *p*= 0.00285, respectively); the multiple adjusted *R^2^* was 0.822, with a *p* value of 4.274e-06. Fig 5 shows the proportion of insects alive after 70 days of the heat treatment for various simulated combinations of temperatures and exposure times.

**Fig. 5.**
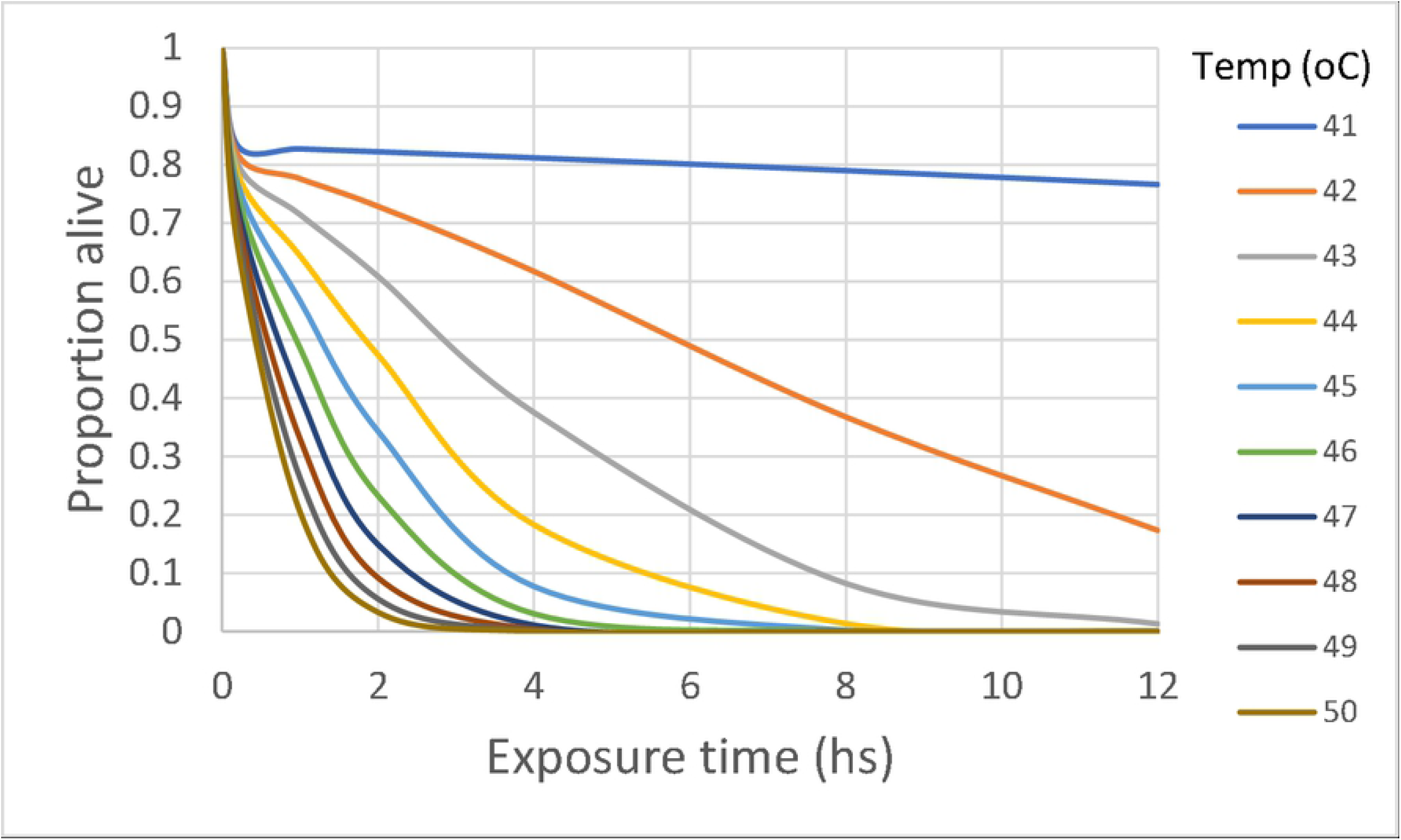
Results of GLM analysis of laboratory experiments. Proportion of insects alive (*l_x_*) after 70 days of the heat treatment for various simulated combinations of temperatures and exposure times.

Another useful parameter to represent the effects of the heat treatments is the expectation of life (e_*x*_) after the day of treatment (*x*); this parameter is defined as the average number of days of life remaining to an individual alive at age or time *x*, and is calculated as 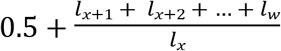, where *l_x_* is the probability of being alive at age or time *x*, and *l_w_* is the same where *w* refers to the time of death [41]. Fig 6 shows the number of days expected to be lived by an average insect after each heat treatment (*e_0_*). These *e_0_* estimates were calculated from both the experimental data and the survival curves predicted by the Cox analysis. It is clearly seen that the variability of the insects’ mortality response becomes more variable as the temperature increases.

**Fig. 6.**
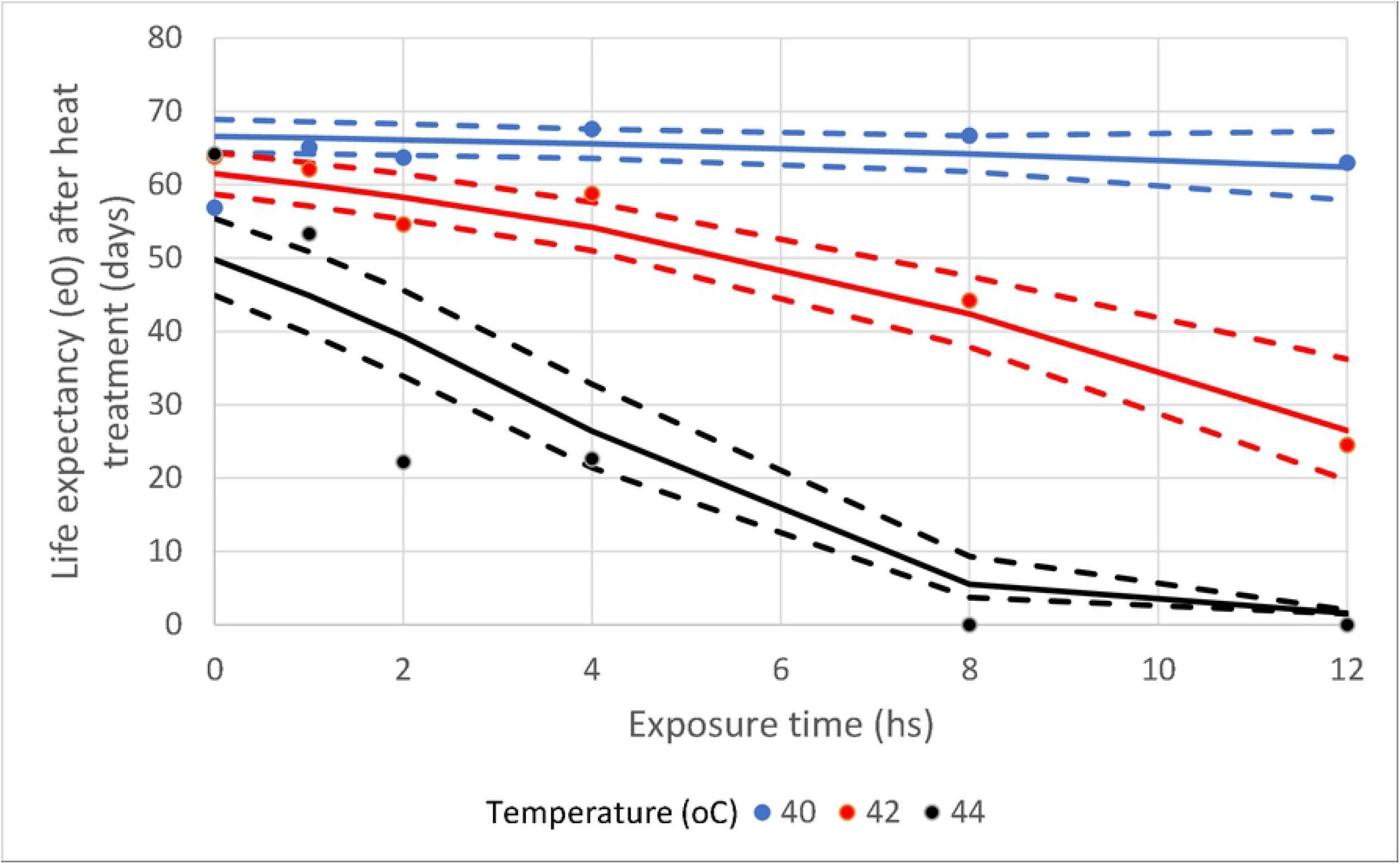
Comparison of observed and expected life expectancies. Life expectancy (*e_0_*, in days) after the day of the heat treatment from both the experimental data and the survival curves predicted by the Cox analysis. The dashed lines are the 95% confidence intervals.

## Discussion

We demonstrate the power of an active learning approach to explore experimental space to design studies that can provide greater understanding of species thermal limits. From a preliminary and very limited dataset, we were able to identify complex but biologically plausible nonlinear interactions between temperature and exposure times shaping mortality, and setting potential thermal limits of triatomine species for SDM (species distribution modelling). This led to successful new experiments that generated novel insights into this important vector group that can be utilized by species distribution models based upon micro-climatic information. This approach can not only guide experimental efforts for the specific study of disease vectors but can also be extended to reduce the experimental envelope in any systems with multiple interacting variables.

The harmful effect on insects’ survival of exposing them to extreme temperatures (close to the upper lethal temperature) for extended periods of time is not linear, and thus difficult to predict [42]. Without our active learning approach, researchers would have had to do a much larger number of experimental combinations to find these non-linear thresholds between exposure and temperature for each species; not only impractical but unaffordable. Our experimental temperatures of between 40-44 °C and exposure times of 1 to 6 hours at high temperatures are found on *T. infestans’* natural habitats during summer [24]; we extended the recorded exposure times to 8 and 12 hours to provide some margin for potential global climate change. Similar to [30], our results showed that survival of fifth nymph of *T. infestans* was not affected by any of the exposure times tested at 40 °C. However, it could affect other physiological traits that were not measured like the germinal cells [43]. In addition, a survival reduction was observed after 8 or more hours of exposure at 42°C or 4 or more hours of exposure at 44°C. Similarly, in nymphs of the related species, *Pastrongylus megystus*, a brief exposure of 1 h to 40°C did not compromise its survival, but a long period of exposure, *i.e.*, 12 h, showed an important drop on survival [27,28]. The combined effect of temperature and exposure time on survival is quite variable, possibly due to thermal tolerance differences across triatomine species, as shown by [44] for seven species of triatomines. Those species were chosen because of their epidemiological relevance, and it was observed that thermo-tolerance range increases with increasing latitude mainly due to better cold tolerances, and limiting their southern distribution [44].

A sub-lethal temperature acquires a considerable ecological relevance, when an extended time of exposures to such a temperature turns into lethal [42]. In our experiments this was the case for 42 and 44 °C. These temperatures did not reduce insects’ survival when exposures times were small, *e.g*., 1 h, while longer exposure times, *i.e*., 2 or 8 h at 44 or 42°C respectively, highly decreases survival. There is little information about the effects of exposure times and temperatures in kissing bugs. There is one study where a brief exposure (0.17 h) at a high temperature (55 °C) produced 100 % mortality in nymphs and adults of *T. infestans*, while lower temperatures presented various effects: at 40 °C there was an increase of insect’s activity, while 50 °C produced severe damage, like knock-down, proboscis extension and leg paralysis [31]. Although the exposure times of our experiments were longer, we observed similar damage, especially knock-down, for almost all exposure times at 44°C, also in *T. infestans* (A.A.C. and I.M. personal observations).

Lastly, our methodology opens up further opportunities to model population dynamics based upon the responses to micro-climatic environments (as [45] have done for in a tephritid fly *Bactrocera dorsalis*).

The effects of temperature and exposure on other demographic parameters like development time and fecundity in order to estimate the thermal effects on the population growth rate are reasonably understood. Under laboratory conditions [46] found in the triatomine *Rhodnius prolixus* that daily temperature fluctuations (DTF) did not affect development time and fertility. However, fecundity was lower in females reared at DTF than at constant temperature, and males had higher body mass reduction rate and lower survival in the DTF regime, suggesting higher costs associated to fluctuating thermal environments [46]. Also, the humidity factor has to be considered when performing SDM, which interacts with temperature (expressed as the vapor pressure saturation deficit [47]). Using an active learning approach could further help to understand this interaction and thus, the dryness dimension of the fundamental niche of small ectotherms such as insects, because of their high surface area-to-volume ratios, are usually at risk of dehydration in arid environments. Undoubtedly the lower thermal tolerance has also to be taken into account for a complete bioclimatic analysis of ectotherm population dynamics. The ultimate goal of all these physiological and SDM modelling efforts is to predict population density in different areas of the triatomine’s geographical range; population density of pathogens’ vectors is an important component of transmission risk, so this kind of micro-SDM modelling is of important epidemiological significance. We hope that our successful results will prompt experiments with other species in that direction.

Incorporating computational advances and active learning into ecological experimental design decision making is rare, but this study provides a case study on how this approach can be used in the context of species thermal ecology. As the world climate shifts even more swiftly, being able to search rapidly through an experimental space to design better experiments to identify species tolerance thresholds is increasingly important. Moreover, as ecological systems are inherently complex and ecological experiments often costly, an active learning approach has the potential to be of more general use in various ecological sub-disciplines.

## Acknowledgements

We are grateful to Yuedong Wang, University of California, Santa Barbara, who made valuable suggestions for the implementation of the GLM analysis to our laboratory results. Raúl Stariolo, of the National Chagas Control Service (Córdoba, Argentina), kindly provided the live specimens of *T. infestans* used to carry out the experiments.

## Authors’ contributions

JR and NFJ conceived the ideas and designed the methodology; PES, AAC and IM designed and carried out the laboratory experiments; all five authors analyzed the data; JR and NFJ led the writing of the manuscript. All five authors contributed critically to the drafts and gave final approval for publication.

## Data availability

The original data and the scripts for running the analyses are hosted at a github site: https://github.com/nfj1380/ThermalLimits_ActiveLearning

